# Different hippocampal subfield volumes predict source memory performance and general cognitive ability in an adult lifespan sample

**DOI:** 10.64898/2026.09.25.754488

**Authors:** Anjana S. Anandan, Marianne de Chastelaine, Ambereen Kidwai, Mingzhu Hou, Claire McIntyre, Michael D. Rugg

## Abstract

Modest positive associations between episodic memory performance and whole hippocampal and hippocampal subfield volumes have been reported in numerous prior studies. A smaller number of studies have reported associations between hippocampal volume and performance on tests of non-mnemonic cognition. The present study examined whether these associations were evident in a lifespan sample of cognitively healthy adults. Of particular interest was whether any identified associations were sensitive to age, and whether associations between subfield volumes and mnemonic and non-mnemonic performance were subfield dependent. We acquired high-resolution T1- and T2-weighted structural images from 163 adults (18-87 years of age). Participants also undertook a comprehensive neuropsychological test battery and an in-scanner test of source memory. Principal components analysis was employed to reduce the neuropsychological test scores to 5 cognitive components. Two components reflected memory performance while the other three reflected different aspects of non-mnemonic cognition. Hippocampal subfields (Cornu Ammonis (CA)1, CA2-3, dentate gyrus (DG) and subiculum) were segmented and measured with the Automated Segmentation of Hippocampus Subfields (ASHS) package. Source memory performance was selectively associated across participants with CA2-3 volume. By contrast, both mnemonic and non-mnemonic component scores derived from the test battery were associated exclusively with the volume of the DG. All associations were age-invariant. The findings indicate that different cognitive domains can be dissociated by virtue of their associations with different hippocampal subfields. Of importance, these associations appear to be life-long and hence are unlikely to reflect individual differences in age-related decline in structural integrity.

## Introduction

Across-participant associations between performance on episodic memory tests and hippocampal volume have been reported in numerous studies. These associations, which typically are weakly positive, have been identified across most of the human lifespan (e.g., Aribisala et al., 2014; Botdorf et al.,2022; Henson et al., 2016; O’Shea et al., 2016; Pohlack et al., 2014). One interpretation of these findings is that they indicate that a relatively large hippocampus confers a modest mnemonic advantage regardless of age (although see, for example, Clark et al., 2020, Fjell et al., 2025, and the meta-analysis of Van Petten et al., 2004, for examples of failures to find evidence of such associations). From this perspective, the findings are consistent with the widely held view that the hippocampus supports computations critical for episodic memory encoding and for the storage and retrieval of recently experienced episodes.

As a result of technical and analytic innovations (e.g., the development of semi-automated image analysis software), the past decade or so has seen an increasing number of reports in which associations between memory performance and hippocampal structural metrics were examined at the subfield level rather than at the level of the whole structure (e.g., Aghjayan et al., 2025; Aslaksen et al., 2018; Aumont et al., 2023; Lai and Chang., 2023; Shing et al., 2011; Zammit et al., 2017). Such studies are partially motivated by evidence that the different subfields play distinct mnemonic roles. For example, whereas the dentate gyrus (DG) and Cornu Ammonis (CA)3 subfields have been implicated, respectively, in the pattern separation and pattern completion operations held to be critical for the encoding, storage and retrieval of unique episodes (O’Reilly et al., 2014), it has been proposed that, among other functions, CA1 supports the learning of statistical regularities across multiple episodes (Singh and Schapiro, 2026). As is the case for whole hippocampal volume, the findings from studies examining relationships between subfield volumes and memory performance mainly take the form of modest positive correlations, albeit with little agreement across studies as to which subfield or subfields demonstrate these associations (across-study differences in how subfields were defined and combined are likely a contributing factor to this disagreement).

Understandably, given the prominent role of the hippocampus in the neuroscience of declarative memory, most studies examining associations between hippocampal volume and cognition have focused on long-term memory performance. Some studies, however, have examined associations with performance on cognitive tasks that seemingly impose few or no mnemonic demands, including intelligence sub-tests. In one early study, Amat et al. (2008) reported a sizeable negative correlation between anterior hippocampal volume and full-scale intelligence in a small sample of young and middle-aged adults. In contrast, Raz et al. (2008) and Reuben et al. (2011) reported modest positive associations (albeit, for older participants only in the latter study) between whole hippocampal volume and measures of fluid intelligence. More recently, Zhu et al. (2017) reported positive associations between CA1 and subiculum volume and performance on the Raven’s Advanced Progressive Matrices Test, a canonical test of fluid intelligence. Together, the findings from these and similar studies (e.g., Colom et al., 2013; Oechslin et al., 2013; but see Aribisala et al., 2014 for an example of null findings in respect of general intelligence) suggest that associations between hippocampal structural metrics and cognitive performance extend beyond long-term memory.

Here, we report data from a study that, among other measures, acquired high-resolution structural MRI images of the hippocampus in a lifespan sample of cognitively healthy adults. Participants undertook a comprehensive neuropsychological test battery that included multiple tests of mnemonic and non-mnemonic ability, including some that would be considered measures of ‘fluid’ cognition (e.g., letter and semantic fluency, Raven’s Progressive Matrices).

Participants also undertook a within-scanner test of source memory (a canonical episodic memory test) that assessed the ability to discriminate between words that had been paired at study with either a face or a scene. We examined across-participant associations between the volumes of 4 hippocampal subfields (CA1, CA2-3, DG and subiculum), cognitive constructs derived from the test battery, and item and source memory metrics obtained from the in-scanner memory test. At issue was whether we could replicate prior findings of associations between subfield volumes and mnemonic performance, whether analogous associations were also evident for non-mnemonic cognitive metrics, whether any associations were moderated by age, and whether volumetric associations between mnemonic and non-mnemonic performance involved different or overlapping subfields.

## Methods

### Participants

A total of 163 cognitively healthy adults contributed to the analyses reported below. The sample was drawn from a larger sample of 188 adults who were recruited as part of a multi-modal lifespan study. The present sample comprised all participants in whom the T1- and T2-weighted structural images required for hippocampal segmentation were acquired and for whom quality control assessments of the images and segmentations (see ‘Estimation of hippocampal subfield volumes’ section below) were acceptable. The participants were recruited from UT Dallas and the surrounding metropolitan Dallas communities. The demographic information pertaining to the 163 participants fulfilling the above criteria are summarized in Table 1.

**Table 1.** Demographic information and neuropsychological test scores (mean (SD)) for the young, middle-aged and older age groups.

|  | Young | Middle-Aged | Older |
| --- | --- | --- | --- |
| N | 37 | 43 | 83 |
| Sex (M/F) | 16/21 | 19/24 | 40/43 |
| Age at neuropsychological test (years) | 22.19 (3.03) | 50.79 (3.37) | 71.25 (5.48) |
| Age range (years) | 18-31 | 45-56 | 64-87 |
| Years of Education | 15.35 (1.72) | 17.15 (2.26) | 16.96 (2.43) |
| Mini Mental State Exam (/30) | 28.65 (1.16) | 29.33 (0.89) | 29.37 (0.87) |
| CVLT<sup>a</sup> recall composite (/64) | 51.08 (7.79) | 49.63 (10.10) | 45.54 (9.11) |
| CVLT<sup>a</sup> recognition hits (/16) | 15.51 (0.77) | 15.02 (1.28) | 14.72 (1.30) |
| CVLT<sup>a</sup> recognition FAs (/32) | 0.78 (0.85) | 1.95 (2.06) | 2.10 (2.28) |
| WMS<sup>b</sup> composite (/100) | 55.59 (11.00) | 52.19 (10.82) | 52.92 (9.20) |
| BVMT<sup>c</sup> recall composite (/48) | 42.35 (3.80) | 37.37 (5.35) | 30.23 (8.41) |
| BVMT<sup>c</sup> recognition hits (/6) | 5.95 (0.23) | 5.86 (0.35) | 5.64 (0.62) |
| Letter Fluency test | 42.14 (10.75) | 46.12 (12.14) | 44.87 (14.01) |
| Symbols Digit Modalities Test (/110) | 57.78 (9.14) | 55.47 (7.36) | 48.41 (7.92) |
| Trail A (seconds) | 20.96 (7.26) | 20.35 (5.58) | 28.21 (7.92) |
| Trail B (seconds) | 51.09 (16.27) | 46.60 (21.94) | 64.25 (21.82) |
| Forward/backward digit span (/64) | 17.89 (4.22) | 18.33 (3.91) | 17.52 (3.40) |
| Category Fluency test | 23.03 (5.41) | 24.37 (6.06) | 20.95 (5.48) |
| Raven's Progressive Matrices (/12) | 10.78 (1.57) | 10.56 (1.33) | 8.94 (1.92) |
| TOPF<sup>d</sup> standard score (scaled score) | 111.76 (8.14) | 114.21 (8.40) | 116.06 (8.39) |

a, California Verbal Learning Test; b, Wechsler Logical Memory; c, Brief Visuospatial Memory
Test-Revised; d, Test of Premorbid Functioning.

All participants were right-handed, had normal or corrected-to-normal vision and were fluent in English. They were compensated at the rate of $30 per hour, and up to $30 for travel. The study was approved by the University of Texas at Dallas Institutional Review Board.

Exclusion criteria during initial screening prior to recruitment included a history of neurological or psychiatric disorder, substance abuse, diabetes, and current or recent use of medications affecting the central nervous system. Participants were accepted into the study according to a common set of inclusion and exclusion criteria based on performance on the neuropsychological test battery (see ‘Neuropsychological test battery’ section below) that were intended to minimize the likelihood of including individuals with cognitive impairment arising from neuropathology.

### Neuropsychological test battery

Participants were administered a battery of neuropsychological tests approximately 2 weeks prior to the MRI session. The battery comprised the Mini-Mental State Examination (MMSE), the California Verbal Learning Test-II (CVLT; Delis et al., 2000), Wechsler Logical Memory (Tests 1 and 2; Wechsler, 2009), the Symbol Digit Modalities Test (SDMT, Smith, 1982), Trail Making (Tests A and B; Reitan and Wolfson, 1985), the F-A-S subtest of the Neurosensory Center Comprehensive Evaluation for Aphasia (Spreen and Benton, 1977), the Wechsler Adult Intelligence Scale–Revised (Forward and Backward digit span subtests; Wechsler, 1981), the Category Fluency test (Benton, 1968), the Wechsler Test of Premorbid Functioning (TOPF, Wechsler, 2011), List 1 of Raven’s Progressive Matrices (Raven et al., 2000) and the Brief Visuospatial Memory Test-Revised (BVMT-R, Benedict, 1997). Scores on each test are summarized in Table 1 for each age group. Participants were excluded from the MRI session if they performed > 1.5 SD below the age-appropriate norms on ≥ 2 non-memory tests, 1 or more memory tests, or if they returned a score of < 26 on the MMSE.

As the 4 CVLT recall scores were highly correlated across participants (min r = 0.75), they were summed to produce a single composite CVLT recall score that was employed for all further analyses. Similarly, the 2 WMS Logical Memory Scores (r = 0.82) and the BVMT recall scores (r = 0.87) were summed to give a single composite score in each case. The correlations between the two Trails scores (A and B) were markedly lower (r = 0.49) and, therefore, they were not aggregated prior to further analysis.

### Scanned memory test

Structural and functional data were acquired during a single MRI scanning session. The experimental memory test comprised two study-test cycles and was undertaken while functional MRI data were acquired (the functional findings will be reported separately). In each study-test cycle participants completed a relational encoding task comprising 96 visually presented critical word-picture pairs (48 word-face pairs, 48 word-scene pairs). During the subsequent retrieval task, 144 words (96 studied or ‘old’ words and 48 unstudied or ‘new’ words) were visually presented. The requirement was to make an old/new recognition judgment for each word and, if the word was judged ‘old’, to indicate whether the word had been paired with a face or a scene at study, or to give a ‘don’t know’ response if the associated item could not be recalled. The duration of each study block was approximately 14 mins, and each test block lasted approximately 18 mins.

Data were aggregated across the two study-test cycles. Estimates of item recognition accuracy (Pr) and source memory accuracy (pSR, i.e., probability of source recollection) were computed for each participant. Pr was estimated as the difference between the overall hit rate and false alarm rate:

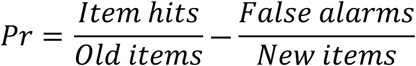

Source memory accuracy (pSR) was computed using only item hits and a modified single high-threshold model (Snodgrass and Corwin, 1988) that accounts for the residual guessing rate (see, for example, Mattson et al., 2014):

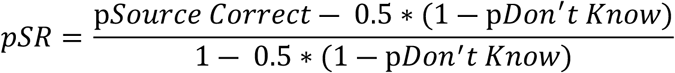

### Principal components analysis

All test scores other than those from the MMSE were subjected to principal components analysis (PCA) to identify the constructs underpinning performance on the test battery (see de Chastelaine et al., 2023, for a similar approach). The PCA was conducted on the full sample of 188 participants (see the Participants sections above). The MMSE was excluded because of ceiling effects, with almost 80% of the sample scoring at (30) or near (29) ceiling. Prior to conducting the PCA, the test scores for each age group were z-transformed within-group to obtain scores that reflected relative test performance independently of age group. The resulting z-scores were combined into a single matrix and subjected to a PCA as implemented in IBM SPSS Statistics v.31. Components were retained if their eigenvalues exceeded 1 (see Table 2). The retained components were subjected to varimax rotation to simplify the solution space. Participant-specific component scores were then computed to reflect the performance of each participant on the retained principal components. This was accomplished by z-scoring the raw test scores across age groups and, for each of the retained components, multiplying the z-scored data by the component loading and summing the resulting scores.

**Table 2.**
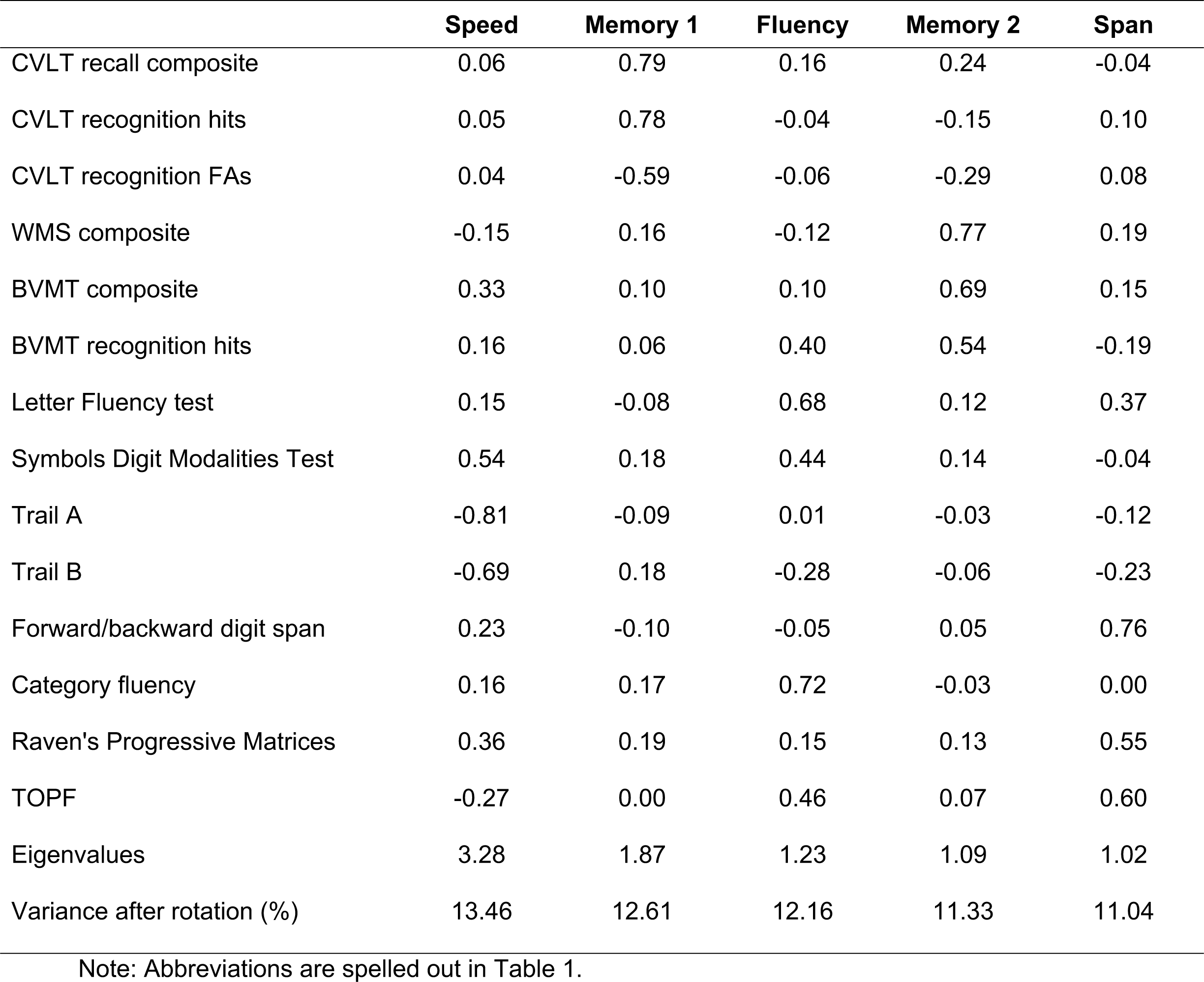
Varimax-rotated factor loadings from the PCA of the neuropsychological test battery.

|  | **Speed** | **Memory 1** | **Fluency** | **Memory 2** | **Span** |
| --- | --- | --- | --- | --- | --- |
| CVLT recall composite | 0.06 | 0.79 | 0.16 | 0.24 | -0.04 |
| CVLT recognition hits | 0.05 | 0.78 | -0.04 | -0.15 | 0.10 |
| CVLT recognition FAs | 0.04 | -0.59 | -0.06 | -0.29 | 0.08 |
| WMS composite | -0.15 | 0.16 | -0.12 | 0.77 | 0.19 |
| BVMT composite | 0.33 | 0.10 | 0.10 | 0.69 | 0.15 |
| BVMT recognition hits | 0.16 | 0.06 | 0.40 | 0.54 | -0.19 |
| Letter Fluency test | 0.15 | -0.08 | 0.68 | 0.12 | 0.37 |
| Symbols Digit Modalities Test | 0.54 | 0.18 | 0.44 | 0.14 | -0.04 |
| Trail A | -0.81 | -0.09 | 0.01 | -0.03 | -0.12 |
| Trail B | -0.69 | 0.18 | -0.28 | -0.06 | -0.23 |
| Forward/backward digit span | 0.23 | -0.10 | -0.05 | 0.05 | 0.76 |
| Category fluency | 0.16 | 0.17 | 0.72 | -0.03 | 0.00 |
| Raven's Progressive Matrices | 0.36 | 0.19 | 0.15 | 0.13 | 0.55 |
| TOPF | -0.27 | 0.00 | 0.46 | 0.07 | 0.60 |
| Eigenvalues | 3.28 | 1.87 | 1.23 | 1.09 | 1.02 |
| Variance after rotation (%) | 13.46 | 12.61 | 12.16 | 11.33 | 11.04 |

Note: Abbreviations are spelled out in Table 1.

### MRI acquisition

Structural MRI data were acquired at the Sammons BrainHealth Imaging Center, University of Texas at Dallas, using a Siemens Prisma 3T scanner equipped with a 32-channel head coil. A T1-weighted anatomical image was acquired with a 3D MPRAGE pulse sequence (FOV = 256 x 256, voxel size 1 x 1 x 1 mm, 160 slices, sagittal acquisition). A high-resolution T2-weighted anatomical image (voxel size 0.4 x 0.4 x 2.0 mm, 52 slices), was also acquired with an oblique coronal orientation perpendicular to the long axis of the hippocampus.

### Estimation of hippocampal subfield volumes

Hippocampal subfields were segmented using the high-resolution T2-weighted images and the Automated Segmentation of Hippocampus Subfields (ASHS; Yushkevich et al., 2015a) software package, in combination with the Princeton Young Adult 3T ASHS Atlas (Hindy et al., 2016). The software and atlas were downloaded from the NITRC repository (https://www.nitrc.org/frs/?group_id=370&release_id=9417) accessed via the ASHS documentation website (https://sites.google.com/view/ashs-dox/home; see Yushkevich et al. (2015a) for a description of the ASHS core algorithms and segmentation pipeline). The hippocampus was segmented into the following subfields: CA1, CA 2-3, DG and subiculum, as shown in Figure 1. As in several prior studies examining hippocampal subfield volumes (e.g., Clark et al., 2023; Iglesias et al., 2015; Winterburn et al., 2013; Yushkevich et al., 2009; see also Yushkevich et al., 2015b) we employed a segregation approach that combined CA2 and CA3 because the two subfields are small and difficult to distinguish reliably on 3T images. Subfields were traced along the entire length of the hippocampus except for the 2-3 slices covering the most anterior portion of the head and the most posterior portion of the tail, regions where there is currently a lack of consensus about subfield boundaries.

**Fig. 1.**
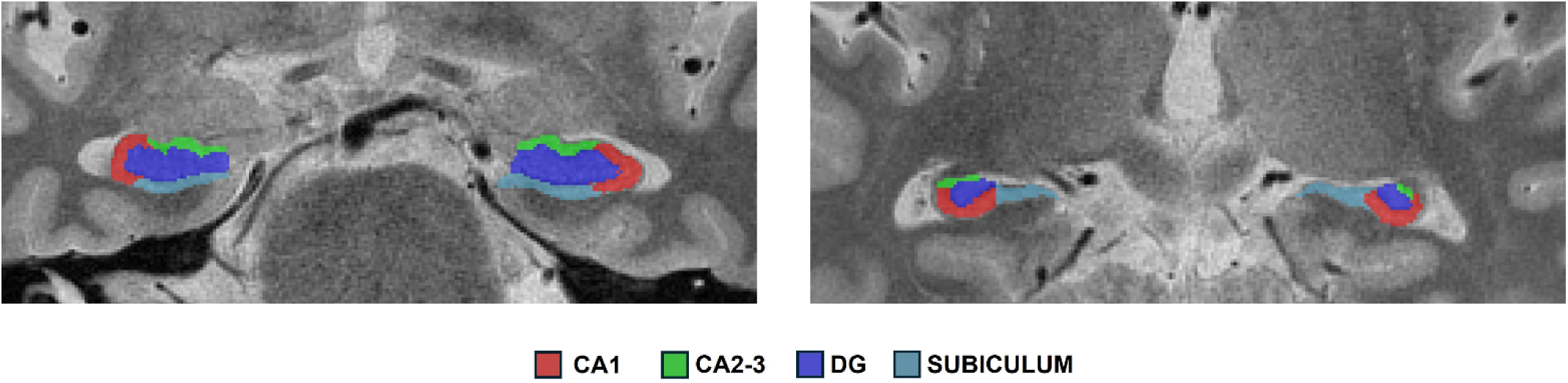
Example of anterior (left) and posterior (right) hippocampi segmented into 4 different subfields by ASHS on a single participant’s T2-weighted image.

Quality control was conducted by 5 of the current authors and was based on group consensus according to the protocols described by Canada et al. (2024). Structural images were rated for artefacts, and landmark visibility on T2 images was also assessed. Segmentation errors were identified and manually edited using ITK-Snap v.4.4.0. Participant-wise estimates of subfield volumes were then extracted. The estimates were residualized on intracranial volume (ICV) prior to further analysis.

### Statistical analysis

Preliminary analyses failed to identify a moderating effect of hemisphere on any association between subfield volumes and cognitive performance. Hence, the volumetric data were collapsed across hemispheres for the purposes of the present analyses.

Age was treated as a continuous variable in the analyses reported below. The effects of age on cognitive performance and hippocampal subfield volumes were evaluated with multiple regression models that employed as predictor variables the linear and quadratic effects of mean-centered age, along with sex and years of education (these latter two variables were included because, either jointly or singly, they were significant predictors of some of the cognitive metrics, see Table 4). To facilitate comparison with prior reports, analyses of the cognitive and volumetric data by age group are reported in Supplemental Results (Supplementary Tables 1 and 2).

Associations between subfield volumes and cognitive performance were examined with two-step hierarchical multiple regression models. The first step in each model employed sex, years of education, and linear and quadratic age terms as predictors. The second step included the relevant subfield volume and its interactions with linear and quadratic age. When the interaction terms were non-significant, the model was re-run after dropping the terms.

Nominal statistical significance was set at p < .05. For the multiple regression analyses examining associations between subfield volume and cognitive performance (which comprised a family of 8 models), statistical significance was Bonferroni-corrected for multiple comparisons; thus, predictor variables were deemed significant at p < .006.

## Results

### Neuropsychological test scores

Demographic information and scores on the tests comprising the neuropsychological test battery are shown in Table 1 segregated by age group.

### Principal components analysis

The varimax-rotated PCA of the neuropsychological test scores returned 5 components that accounted for over 60% of the variance of the scores. The loadings of the different tests along with the associated eigenvalues and proportions of variance explained are reported in Table 2. The components were named according to the tests with the highest loading on each component, namely: ‘speed’, ‘fluency’, ‘span’, and two components (‘memory 1’ and ‘memory 2’) that reflected performance on the different long-term memory tests included in the battery.

The 5 component scores, along with their average (henceforth, general cognitive ability) are reported in Table 3, segregated by age group. Also included in the table are the 2 metrics derived from the experimental memory test (see ‘Scanned memory test’ above), item (Pr) and source (pSR) memory performance. The outcomes of the regression models examining the effects of age on these behavioral metrics are summarized in Table 4 and illustrated in Figure 2. As is evident from the table, linear age effects were reliable in each case, reflecting a monotonic decline in source memory performance with increasing age. However, in the case of speed, fluency, span and general cognitive ability, linear effects were accompanied by quadratic effects indicating, for those tests, that cognitive ability demonstrated an accelerating decline with increasing age.

**Fig. 2.**
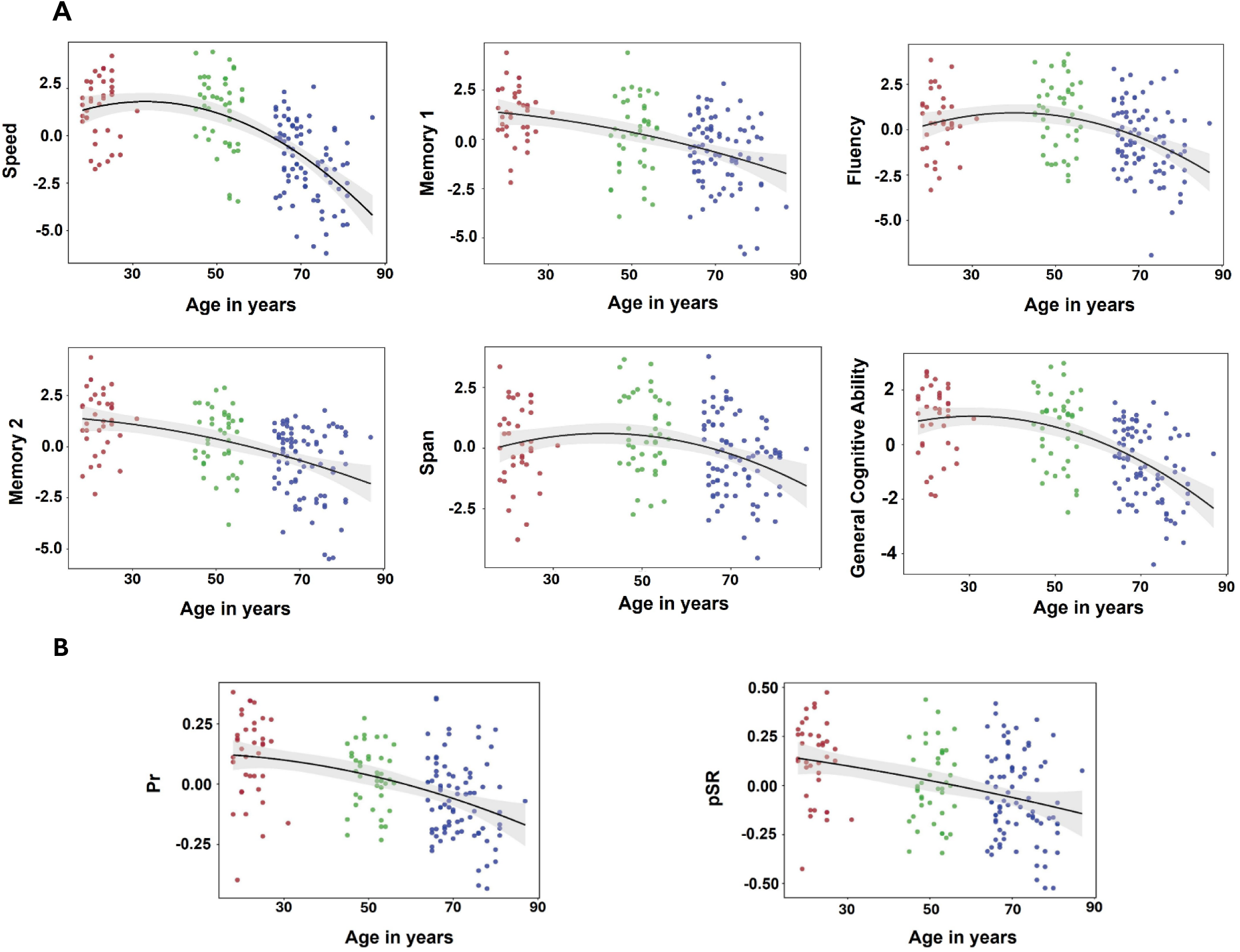
Scatterplots depicting the relationships between A) Cognitive component scores (residualized on years of education and sex) and age; B) Item and source memory scores (residualized on years of education and sex) and age.

**Table 3.** Individual component, general cognitive ability (Gen Cog) and in-scanner memory test scores (mean (SD)) segregated by age group.

|  | Young | Middle-Aged | Older |
| --- | --- | --- | --- |
| Speed | 1.44 (1.66) | 1.17 (1.94) | -1.51 (1.98) |
| Memory 1 | 1.25 (1.21) | 0.26 (2.10) | -0.73 (1.96) |
| Fluency | 0.25 (1.59) | 0.81 (1.90) | -0.63 (1.98) |
| Memory 2 | 1.26 (1.43) | 0.41 (1.59) | -0.74 (1.77) |
| Span | -0.05 (1.65) | 0.56 (1.62) | -0.35 (1.65) |
| Gen Cog | 0.83 (1.19) | 0.64 (1.38) | -0.79 (1.37) |
| Pr | 0.64 (0.17) | 0.56 (0.12) | 0.46 (0.17) |
| pSR | 0.65 (0.20) | 0.52 (0.19) | 0.45 (0.22) |

### Associations between age and hippocampal subfield volumes

The volumetric data (residualized by ICV, sex and years of education) are summarized by age group in Table 5. The outcomes of the regression models examining the effects of age are summarized in Table 6. As is evident from the table, significant linear and quadratic age effects were identified for CA1 and the subiculum. These effects reflected a tendency for the volumes to peak around middle age and to decline thereafter, as depicted in Figure 3.

**Fig. 3.**
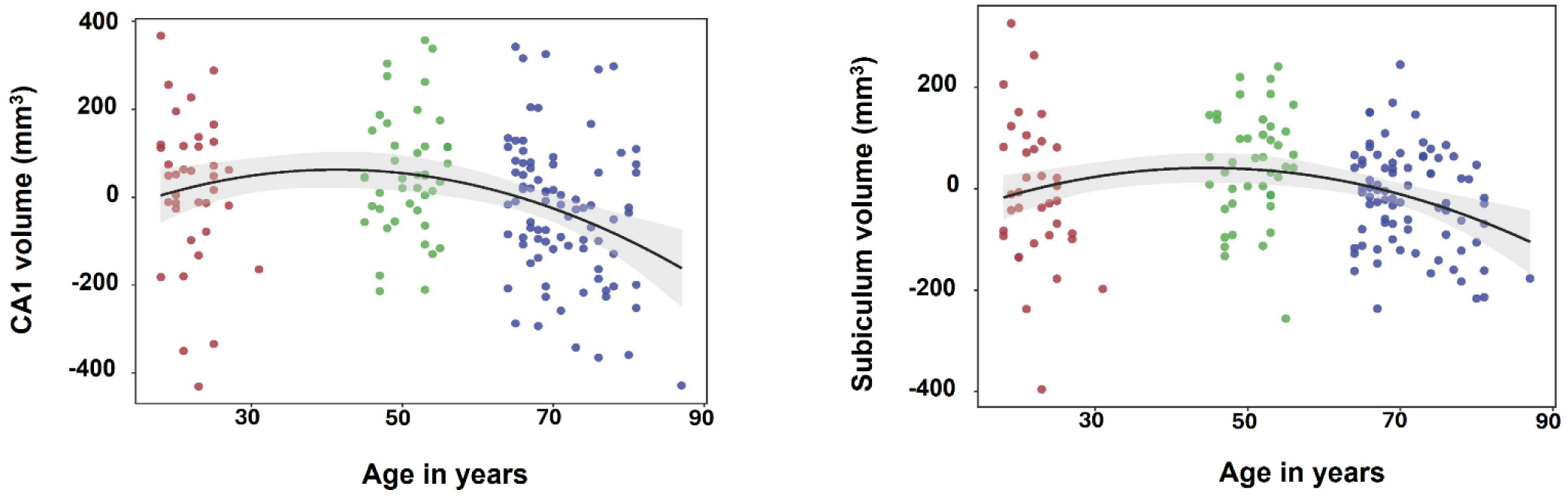
Scatterplots depicting the associations between age and CA1 (left) and subiculum (right) volumes residualized on ICV, years of education and sex.

### Associations between cognitive performance and hippocampal subfield volumes

As described in Methods, we employed hierarchical regression to examine associations between cognitive performance, age and subfield volumes. Preliminary analyses failed to identify evidence for unique associations between any individual cognitive construct and subfield volumes, as evidenced by the finding that no association attained the corrected significance level after the mean of the remaining component scores was entered in the model as an additional covariate (see Hou et al, 2021, for a similar approach to identifying unique associations). Accordingly, we employed a single cognitive construct – general cognitive ability – to represent performance on the neuropsychological test battery. For analogous reasons, we selected pSR as the metric representative of performance on the experimental memory test.

We employed a set of 8 regression models in the primary analyses reported below. Four of these examined associations between the different subfields and pSR, while the other 4 examined associations with general cognitive ability. The first step of each model comprised the same set of predictor variables, namely, sex, years of education, and linear and quadratic age terms. In the second step the relevant subfield volume and its interaction terms with linear and quadratic age were added. In all analyses other than that examining associations between general cognitive ability and CA2-3 and subiculum volumes, the interaction terms were non-significant and dropped from the final model.

Since the first step of each regression model is identical to the models reported in Table 4, we focus on whether the inclusion in the second step of the volumetric predictors led to a statistically significant increase in explained variance. The results of these analyses are reported in Tables 7 and 8. Turning first to pSR, it can be seen from Table 7 that inclusion of CA2-3 volume led to a significant increase in the explained variance of pSR (partial r = .249 for the association between pSR and CA2-3 volume), while subiculum volume failed to survive multiple correction, and DG and CA1 volumes were non-significant predictors. The association between CA2-3 volume and pSR is illustrated in Figure 4. Table 8 summarizes the outcomes of the addition to the model of the volumetric predictors for general cognitive ability. As can be seen from the table, after correction for multiple comparisons, the only subfield that led to a significant increase in explained variance was DG (partial r = .246 for the association between general ability and DG volume). This association is illustrated in Figure 5.

**Table 4.** Multiple regression results for sex, years of education and age predicting cognitive performance (p values in parentheses).

| | Standardized $\beta$ | t (p) | F (p) | R<sup>2</sup> |
| --- | --- | --- | --- | --- |
| <u>Speed</u> |  |  | 27.53 (<.001) | 0.41 |
| Intercept |  | -0.17 (.862) |  |  |
| Sex | 0.00 | 0.01 (.991) |  |  |
| Years of education | 0.05 | 0.76 (.448) |  |  |
| Linear age | -0.80 | -10.24 (<.001) |  |  |
| Quadratic Age | -0.34 | -4.43 (<.001) |  |  |
| <u>Memory 1</u> |  |  | 18.13 (<.001) | 0.32 |
| Intercept |  | -2.10 (.037) |  |  |
| Sex | -0.34 | -5.16 (<.001) |  |  |
| Years of education | 0.20 | 2.89 (.004) |  |  |
| Linear age | -0.49 | -5.86 (<.001) |  |  |
| Quadratic Age | -0.06 | -0.66 (.507) |  |  |
| <u>Fluency</u> |  |  | 6.68 (<.001) | 0.15 |
| Intercept |  | -1.05 (.293) |  |  |
| Sex | -0.06 | -0.82 (.414) |  |  |
| Years of education | 0.13 | 1.68 (.095) |  |  |
| Linear age | -0.46 | -4.94 (<.001) |  |  |
| Quadratic Age | -0.31 | -3.35 (.001) |  |  |
| <u>Memory 2</u> |  |  | 15.32 (<.001) | 0.28 |
| Intercept |  | -2.20 (.029) |  |  |
| Sex | -0.18 | -2.68 (.008) |  |  |
| Years of education | 0.20 | 2.76 (.007) |  |  |
| Linear age | -0.55 | -6.44 (<.001) |  |  |
| Quadratic Age | -0.07 | -0.84 (.404) |  |  |
| <u>Span</u> |  |  | 5.14 (<.001) | 0.12 |
| Intercept |  | -2.25 (.026) |  |  |
| Sex | 0.08 | 1.08 (.284) |  |  |
| Years of education | 0.21 | 2.58 (.011) |  |  |
| Linear age | -0.34 | -3.61 (<.001) |  |  |
| Quadratic Age | -0.25 | -2.68 (.008) |  |  |
| <u>General Cognitive Ability</u> |  |  | 20.41 (<.001) | 0.34 |
| Intercept |  | -2.05 (.042) |  |  |
| Sex | -0.13 | -2.02 (.045) |  |  |
| Years of education | 0.19 | 2.84 (.005) |  |  |
| Linear age | -0.70 | -8.53 (<.001) |  |  |
| Quadratic Age | -0.27 | -3.34 (.001) |  |  |
| <u>Pr</u> |  |  | 11.65 (<.001) | 0.23 |
| Intercept |  | 3.57 (<.001) |  |  |
| Sex | -0.05 | -0.75 (.454) |  |  |
| Years of education | 0.17 | 2.30 (.023) |  |  |
| Linear age | -0.54 | -6.03 (<.001) |  |  |
| Quadratic Age | -0.08 | -0.94 (.348) |  |  |
| <u>pSR</u> |  |  | 6.82 (<.001) | 0.15 |
| Intercept |  | 3.42 (<.001) |  |  |
| Sex | 0.00 | -0.00 (.997) |  |  |
| Years of education | 0.05 | 0.63 (.531) |  |  |
| Linear age | -0.40 | -4.28 (<.001) |  |  |
| Quadratic Age | -0.01 | -0.10 (.918) |  |  |

**Table 5.** Summary of subfield volume data (mean (SD)) residualized on ICV, sex and years of education, segregated by age group.

| Subfield volumes | Young | Middle-Aged | Older |
| --- | --- | --- | --- |
| CA1 | 18.36 (170.90) | 51.83 (137.30) | -35.03 (161.80) |
| CA2-3 | -23.23 (103.50) | 0.02 (91.96) | 10.35 (96.79) |
| DG | 7.62 (207.78) | 33.31 (187.76) | -20.65 (228.72) |
| Subiculum | -7.81 (140.00) | 45.98 (106.63) | -20.34 (96.53) |

**Table 6.**
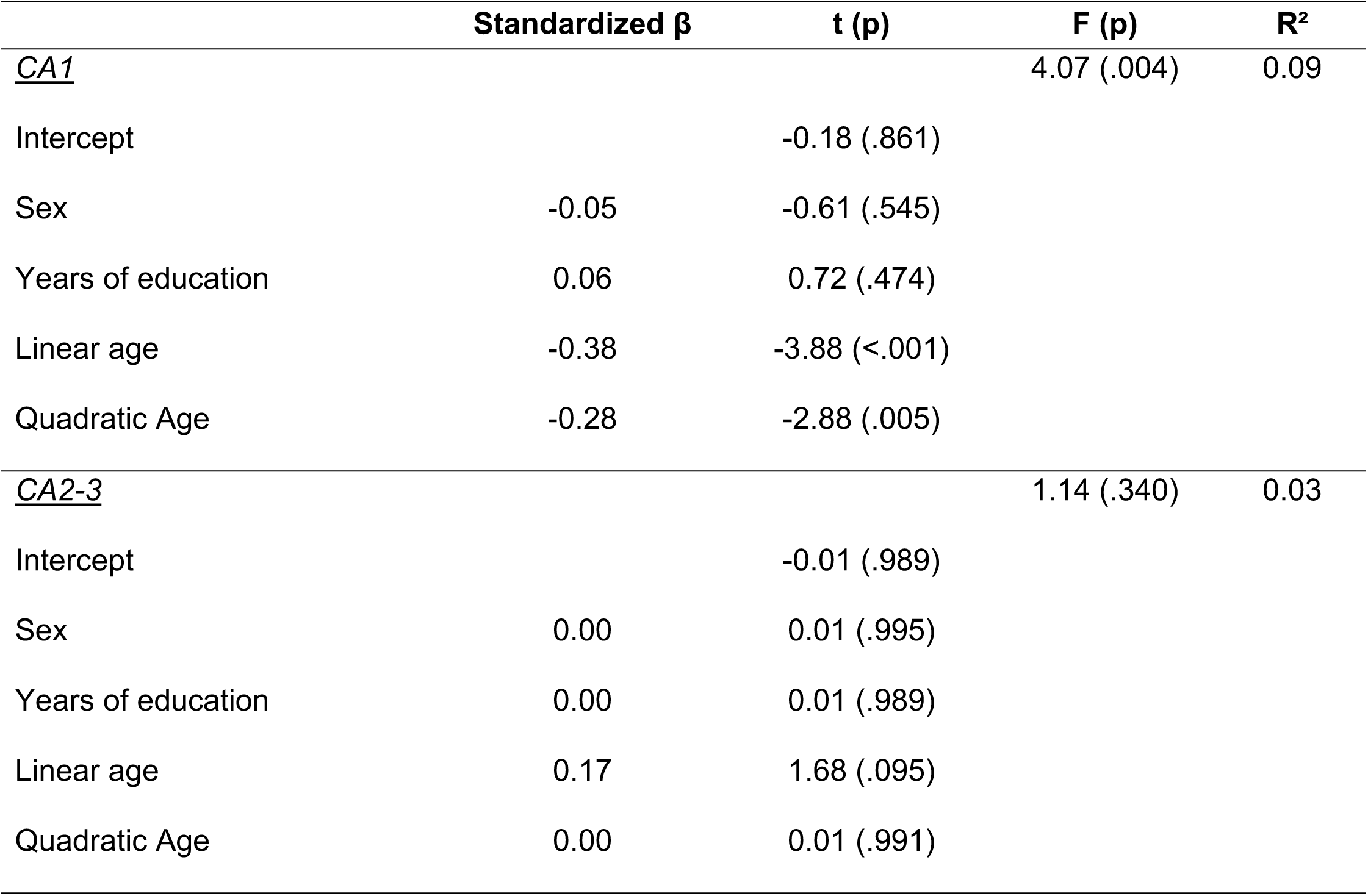
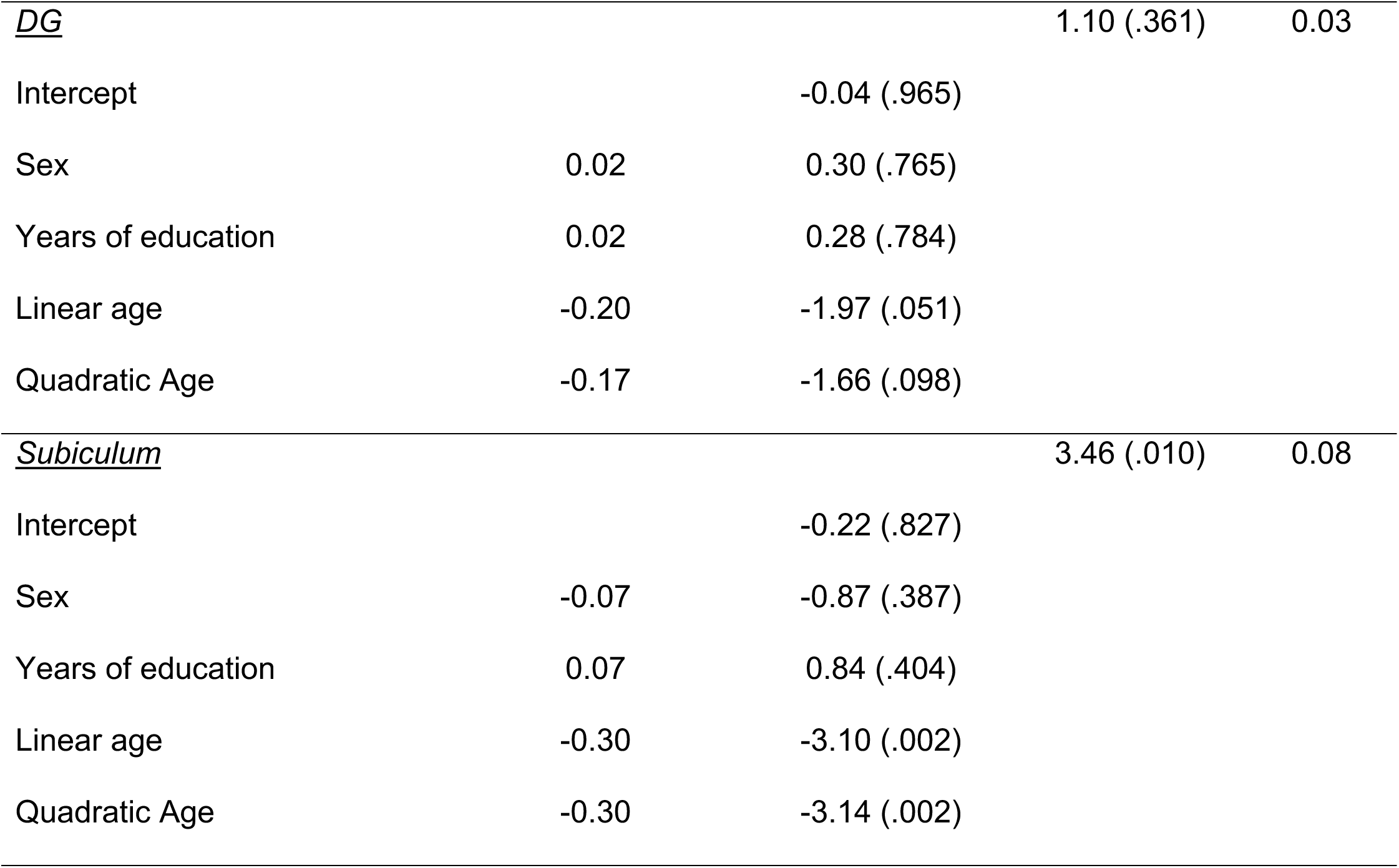
Multiple regression model results for sex, years of education and age predicting hippocampal subfield volumes.

**Fig 4.**
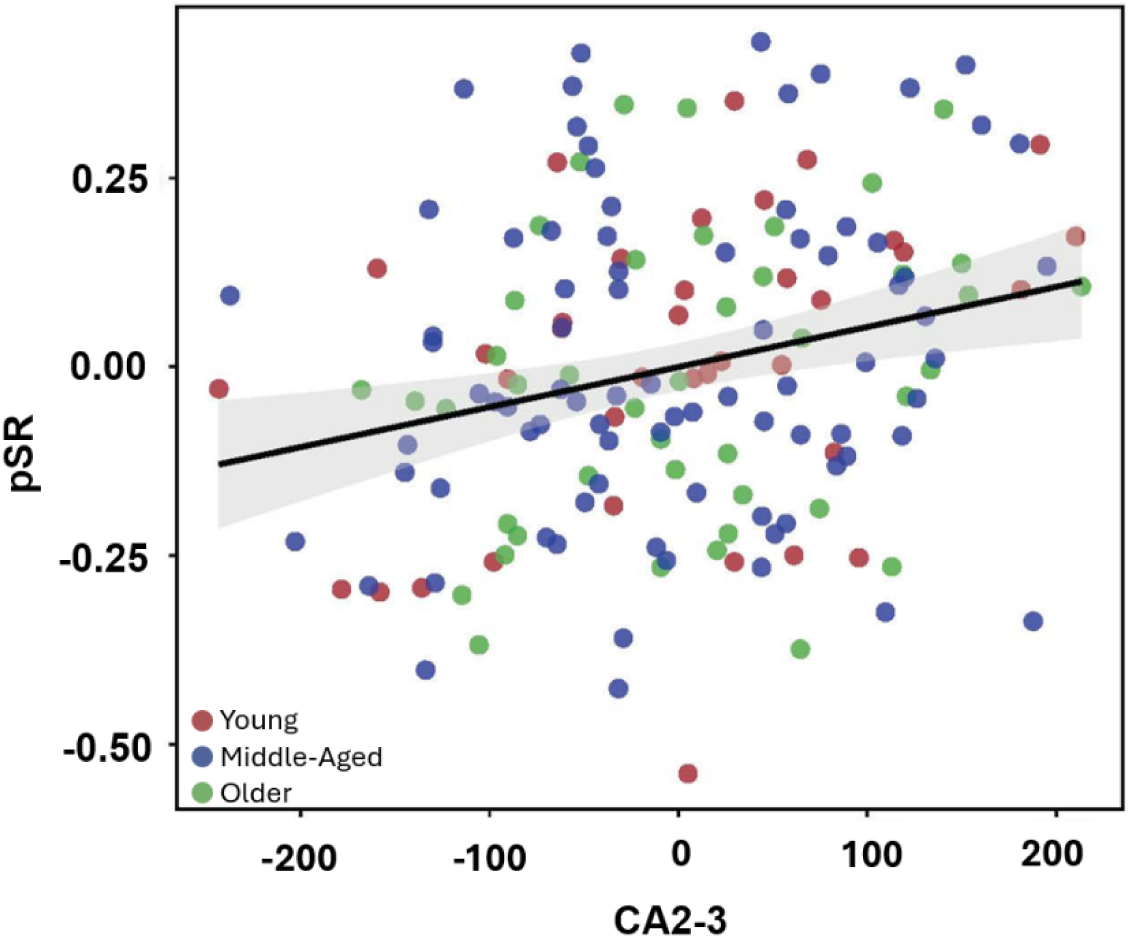
Scatterplot depicting the association between CA2-3 volume (after controlling for sex, years of education, and the linear and quadratic effects of age) and pSR across all participants.

**Fig 5.**
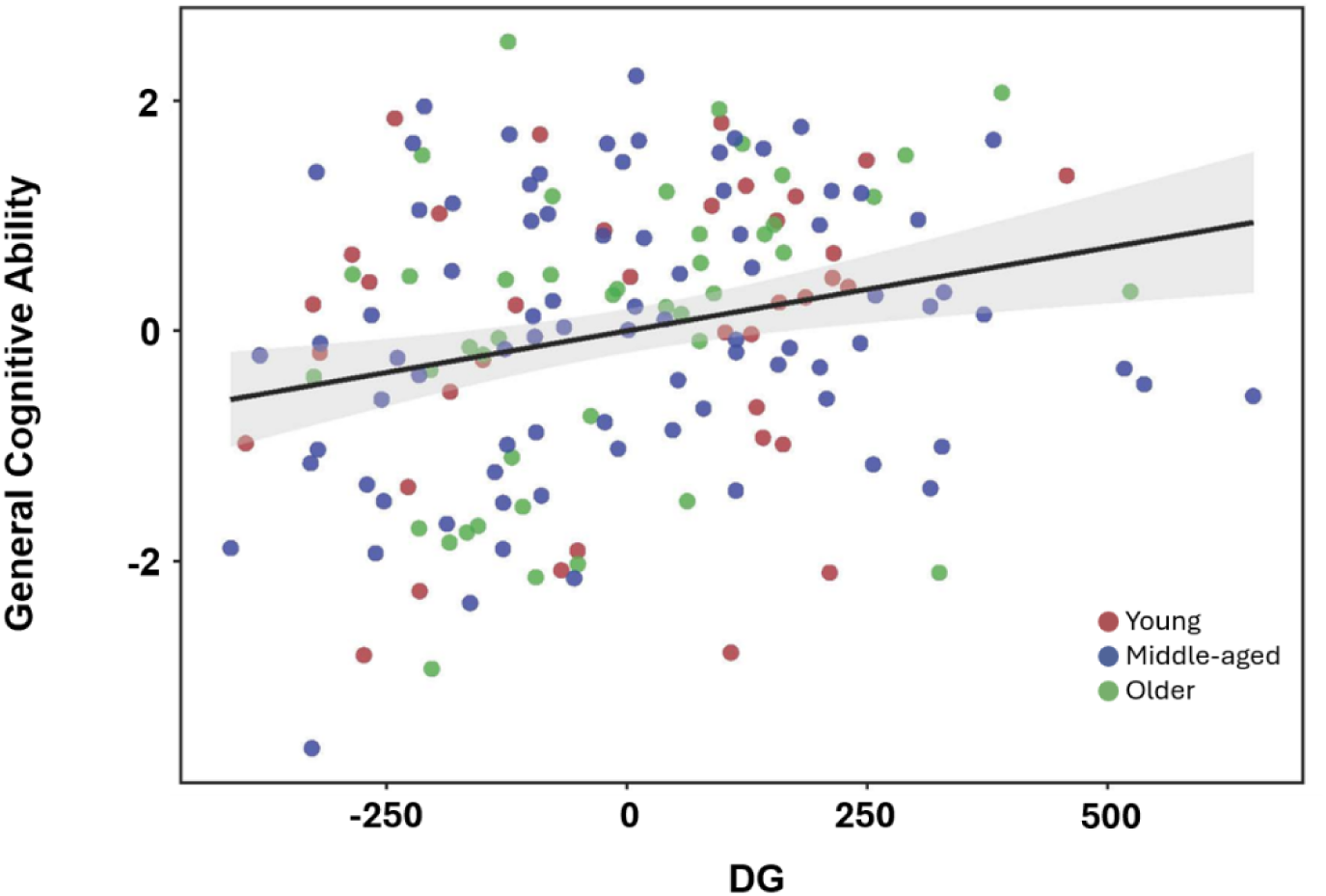
Scatterplot depicting the association between dentate gyrus volume (after controlling for sex, years of education, and the linear and quadratic effects of age) and general cognitive ability across all participants.

**Table 7.** Hierarchical multiple regression results for hippocampal subfield volumes predicting pSR.

| | Standardized $\beta$ | t (p) | F (p) | $\Delta F$ (p) | R<sup>2</sup> | $\Delta R^2$ |
| --- | --- | --- | --- | --- | --- | --- |
| <u>Step 1</u> |  |  | 6.82 (< .001) |  | 0.15 |  |
| Intercept |  | 3.42 (< .001) |  |  |  |  |
| Sex | 0.00 | -0.00 (.997) |  |  |  |  |
| Years of education | 0.05 | 0.63 (.531) |  |  |  |  |
| Linear age | -0.40 | -4.28 (< .001) |  |  |  |  |
| Quadratic Linear age | -0.01 | -0.10 (.918) |  |  |  |  |
| <u>Step 2</u> |  |  | 6.27 (< .001) | 3.59 (.060) | 0.17 | 0.02 |
| CA1 | 0.15 | 1.90 (.060) |  |  |  |  |
| <u>Step 2</u> |  |  | 7.86 (< .001) | 10.38 (.002) | 0.20 | 0.05 |
| CA2-3 | 0.23 | 3.22 (.002) |  |  |  |  |
| <u>Step 2</u> |  |  | 5.86 (< .001) | 1.86 (.175) | 0.16 | 0.01 |
| DG | 0.10 | 1.36 (.175) |  |  |  |  |
| <u>Step 2</u> |  |  | 6.64 (< .001) | 5.17 (.024) | 0.17 | 0.03 |
| Subiculum | 0.17 | 2.27 (.024) |  |  |  |  |

**Table 8.**
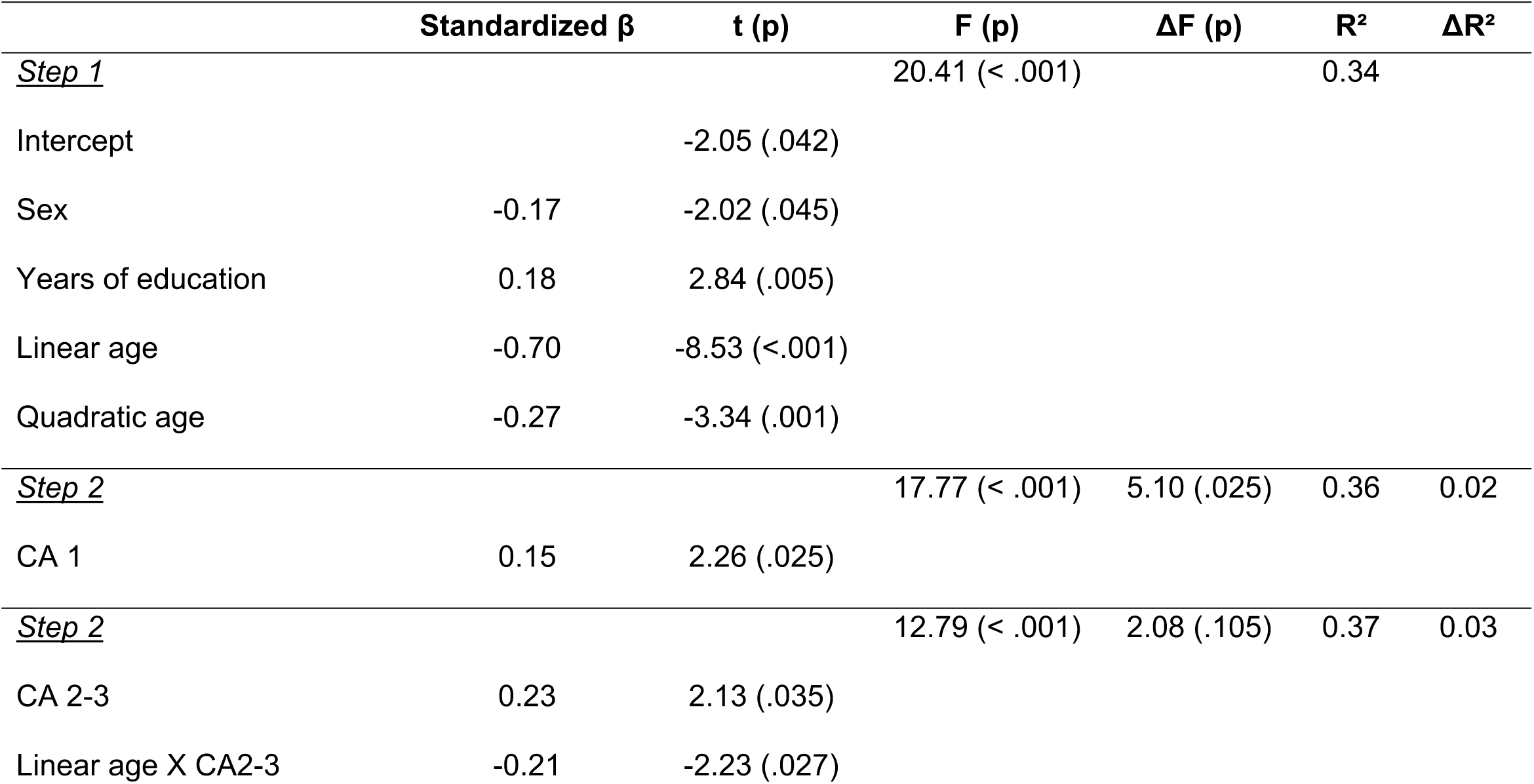
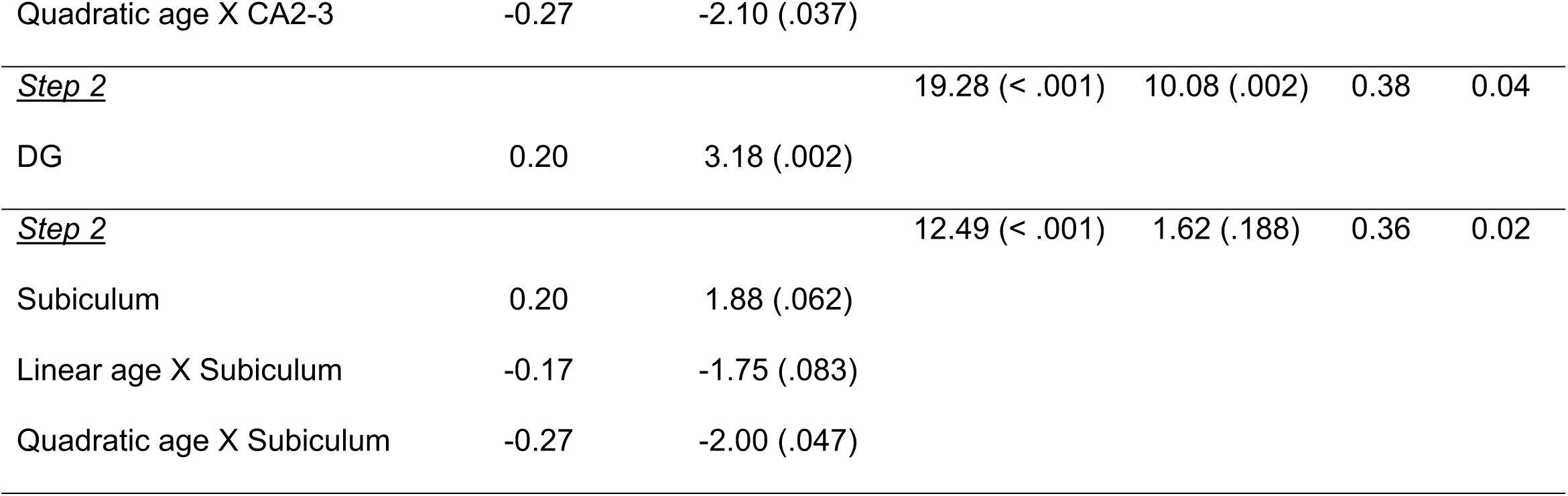
Hierarchical multiple regression results for hippocampal subfield volumes predicting general cognitive ability.

### Specificity of associations between cognitive performance and subfield volumes

We went on to examine the specificity of the associations between pSR and general cognitive ability and, respectively, CA2-3 and DG volumes. First, we constructed a regression model in which, in addition to the predictors of sex, years of education and the two age terms, all 4 subfield volumes were included as predictors (note that the volumes were moderately inter-correlated, the correlations ranging from r = 0.34 to r = 0.65). Despite these collinearities, CA2-3 remained a significant predictor of pSR, as did the DG for general cognitive ability [partial rs = 0.19 (p = .019) and 0.19 (p = .016) respectively]. Second, we ascertained that the two associations remained reliable when the models predicting each performance metric included the other metric as an additional predictor, e.g., predicting pSR with the variables of sex, years of education, the two age terms, CA2-3 volume *and* general cognitive ability. In both cases, the associations remained significant (partial r for pSR and CA2-3 = 0.24, p = .002, partial r for general cognitive ability and DG = 0.22, p = .005).

### Associations between non-mnemonic cognitive components and subfield volumes

As already noted, the general cognitive ability component was employed in the analyses reported above because no single cognitive component accounted for variance in subfield volumes beyond that explained by general ability. However, two of the individual components comprising the composite score reflect mnemonic performance. To examine whether the association reported above between general cognitive ability and DG volume was driven by these components, we repeated the analyses employing a non-mnemonic composite component score comprising the average of the speed, fluency and span scores. This yielded findings closely similar to those reported for the full composite component (see Supplemental Materials, Supplemental Table 3). Notably, the association between the non-mnemonic component score and DG volume (r = 0.23, p = .004, controlling for sex, years of education and linear and quadratic age) remained significant. The analogous correlation for the aggregate of the 2 mnemonic components was r = 0.18, p = .026. The two correlations did not differ significantly.

### Whole hippocampal volume

Although the focus of the present analyses is at the subfield level, we also performed exploratory analyses at the level of the whole hippocampus, allowing comparison with the findings of prior studies that employed this metric. The full analysis can be found in Supplemental Materials, Supplemental Tables 4-6. In brief, after controlling for years of education and sex, total hippocampal volume demonstrated both linear and quadratic effects of age and, after controlling for these variables (along with sex and years of education), it correlated modestly with both pSR and general cognitive ability (r = 0.20, p = .012 and r = 0.21, p = .010, respectively).

## Discussion

The main goal of the present study was to examine associations between the volumes of different hippocampal subfields, age, and performance on a variety of mnemonic and non-mnemonic cognitive tests. We found little evidence that hippocampal subfield volumes explained independent sources of variance across the 5 different cognitive components represented by the neuropsychological test battery, including two components that principally reflected performance on tests of long-term memory. Rather, associations between the cognitive component scores and subfield volumes were fully captured by the association between a general ability score and DG volume. DG volume was not, however, associated with performance on the within-scanner source memory test, which instead correlated exclusively with CA2-3 volume. Notably, both associations were age-invariant.

We first discuss the effect of age on cognitive performance and subfield volumes. In the case of cognitive performance, all scores demonstrated either a linear or a positively accelerating decline with increasing age. These findings are broadly consistent with those reported in numerous prior cross-sectional studies (for review, see Yang et al., 2023). Age effects were also evident for the volumes of CA1 and the subiculum (but not CA2-3 or DG). In both cases, the effects were non-linear, such that the volumes demonstrated a tendency to peak in middle age and to decline thereafter. These findings are seemingly at odds with those from a recent meta-analysis of 30 studies of cognitively healthy adults that examined age effects on subfield volume (Homayouni et al., 2023) and identified a modest negative linear association with age (max r = - .19) for each included subfield (CA1, DG-CA3-4, subiculum). The present findings are, however, characteristic of the diverse findings reported across individual studies (see Homayouni et al., 2023 for review). For example, in contrast to the present findings, Daugherty et al. (2016) reported a linear age effect on CA1-2 volume in a sizeable lifespan sample, along with a quadratic effect on CA3-DG, and a null effect for the subiculum. By contrast, also in an adult lifespan sample, Foster et al. (2019) reported exclusively linear age effects for CA1-2, CA3-DG and the subiculum. And, somewhat reminiscent of the present findings, Gohel et al. (2025) reported quadratic age effects in a composite CA sub-region, DG, and subiculum, with volumes peaking between 30 and 40 years of age. It is unclear to what extent these inconsistent findings across studies reflect differences in substantive variables such as sample composition or choice of subfield combinations, rather than the confounding influence of nuisance variables such as scanner model or image pre-processing and segmentation methods.

Turning to the associations between subfield volumes and cognitive performance, the present findings are broadly consistent with previous reports that total hippocampal volume, and the volumes of individual subfields, are predictive of performance on both tests of memory and cognitive tests that appear to make minimal demands on long-term memory (see Introduction). Arguably, the most salient aspects of the present findings are that the associations identified between subfield volume and cognitive performance were age-invariant, and that in-scanner source memory performance and general cognitive ability scores were associated with distinct subfields.

The finding that the two associations identified between cognitive performance and subfield volume were age-invariant is consistent with the findings of some (e.g., Aslaksen et al., 2018; Henson et al., 2016; Pohlack et al., 2014; Zhu et al., 2017), but not all (e.g., Reuben et al., 2011) of the relatively few studies that have examined such associations in young adult or lifespan samples rather than in later-life samples only. The additive effects of age and subfield volume on cognitive performance suggest that the across participant associations identified between volume and performance reflect co-variance that was stable across the age range.

This suggestion gains further support from the finding that, when the analyses were repeated after excluding the older age group (leaving a sample of 80 participants aged 18 – 56 yrs), both associations remained reliable (CA2-3 and pSR: r = .29, p = .013; DG and general cognitive ability: r = .30, p = .008, in both cases controlling for sex, years of education, and linear and quadratic age effects). Relatedly, the null effects of age on the CA2-3 and DG volumes indicate that, in contrast to some prior findings (e.g., Foster et al., 2019; Gohel et al., 2025), the marked age differences that were evident in cognitive performance were not mediated by the effects of age on subfield volume.

The in-scanner test of source memory employed in the present study belongs to a class of associative memory tests that, according to neuropsychological (Cooper et al., 2017; Yonelinas et al., 1998) and functional neuroimaging (Rugg and Vilberg, 2013) evidence, place substantial demands on the hippocampus. In the present study, source memory performance was positively associated with CA2-3 volume. This finding is partially consistent with prior reports of positive associations between subfield volumes and episodic memory performance, but to our knowledge only one prior study has reported an association selectively with CA2-3 (Clark et al., 2023). In that study, a positive association was identified between CA2-3 volume and the number of ‘internal’ (episodic) event details recalled in an autobiographical memory test administered to a large sample of young adults (intriguingly, it has been reported that localized atrophy of CA3 consequent to limbic encephalitis results in impaired autobiographical memory for internal events; Miller et al., 2017). The association was present only in low performing participants and was not evident for performance on a range of other memory tests, such as paragraph recall. The authors suggested that the association with autobiographical recall reflected the contribution of CA3 (and, particularly, its posterior part) to pattern completion, which they speculated might be especially important for the retrieval of vivid ‘real-world’ memories. The present findings extend those of Clark et al. (2023) to a standard laboratory memory test, specifically, a test that requires the encoding and subsequent retrieval after a few minutes of novel inter-item associations. Not all forms of associative memory correlate with CA2-3 volume, however: using a task requiring associative recognition of Chinese character pairs, Lai and Chang (2023) reported reliable correlations with performance for the volumes of all subfields other than CA2-3, albeit in an analysis that collapsed across young and older age groups but did not control for age, raising the possibility that the reported correlations were confounded by this variable.

Considering the proposed role of CA3 in the storage and retrieval of pattern-separated memory representations (O’Reilly et al., 2014), the present finding that source memory performance was associated with a putative index of the efficacy of these neurocognitive operations is arguably unsurprising. More surprising, perhaps, is the specificity of the association, especially given the important role in the encoding of high-fidelity memories attributed to the DG (Yassa and Stark, 2011). Also notable is that the finding for source memory did not extend to either of the two memory components derived from the neuropsychological test battery, even though performance on their constituent tests (CVLT in the case of memory 1, WMS and BVMT-R for memory 2) is hippocampally dependent (e.g., Rempel-Clower et al., 1996). It is noteworthy, however, that source memory performance and the two memory component scores were only moderately correlated (partial r = .39 and partial r = .34 for pSR and memory 1 and memory 2 respectively), raising the possibility that only source memory placed significant demands on the computations supported by CA2-3. Consistent with this possibility, the association between source memory performance and CA2-3 volume was barely affected by inclusion in the regression model of the two memory component scores as additional covariates (partial r = .242, p =.002).

Turning to the cognitive component scores, as was noted previously, we were unable to identify any reliable associations with subfield volumes that were unique to a specific component. Rather, the individual component scores were encapsulated by a single general cognitive ability score that demonstrated a robust association with DG volume. Of importance, this association was not driven by the memory component scores; in fact, as was noted in the Results, the composite of the three non-mnemonic components (speed, fluency and span) correlated more strongly with DG volume than did the aggregated memory scores.

As reviewed in the Introduction, several prior studies have described positive associations between either whole hippocampal or subfield volume and what might loosely be referred to as ‘fluid’ cognition, including in samples exclusively comprising young adults (e.g., Pohlack et al., 2014), although none, to our knowledge, reported an association as specific as that identified here). Together, the previous and current findings leave little doubt that hippocampal volume is a predictor of non-mnemonic cognitive abilities (notably, those reflected in fluid intelligence), raising the possibility that one or more hippocampal subregions support processes that contribute to these abilities. Of relevance to this possibility, it has been reported that fluid intelligence is weakly correlated with the fidelity of the hippocampal representation of a learned two-dimensional space, as reflected by fMRI multi-voxel pattern analysis (Tenderra and Theves, 2025). Whether this association reflects a direct contribution of hippocampal relational processing to fluid intelligence (the interpretation preferred by Tenderra and Theves), rather than the contribution of processes that operate independently of the hippocampus but whose efficacy correlates with DG volume, is unclear, however. This latter possibility is arguably more compatible with the seeming absence of neuropsychological evidence linking the hippocampus to fluid cognition. Notably, one of the defining characteristics of hippocampal amnesia is a marked disparity between performance on tests of memory and intelligence. For example, in the severely amnesic patient HM, who sustained surgical damage not only to the hippocampus but the surrounding medial temporal neocortex also, intelligence test scores *increased* from pre-to post-surgery (Scoville and Milner, 1957). More generally, so far as we can ascertain, there is little to no published evidence that patients with damage restricted to the hippocampus score lower on intelligence tests (fluid or crystallized) than control participants; indeed, a large lesion-mapping study reported that the principal neural region demonstrating an association with fluid intelligence was right prefrontal cortex (Cipolotti et al., 2023). It is of course possible that sufficiently well-powered studies will reveal an association between hippocampal damage and fluid cognition, but it seems likely that any such association will be subtle and dwarfed by the impact of the lesion on memory performance. In the absence of such evidence, the reason why hippocampal volume should correlate with non-mnemonic cognitive performance remains obscure.

We note that the foregoing discussion is based on the assumption that ‘bigger is better’; that is, a relatively large neural region confers a computational advantage over a smaller one, at least in adulthood. There is however little in the way of neurobiological theory to support this assumption, which gains its support primarily through numerous reports (including the present study) of positive associations between regional size and cognitive performance (but see, for example, Amat et al., 2008 and Colom et al., 2013 for examples of negative associations between adult hippocampal volume and cognition, along with interpretations framed in terms of the greater efficiency of relatively sparse microcircuitry). Additionally, reports of null associations in cognitively healthy adults between hippocampal volume and the magnitude of task-evoked functional activity, and the apparent absence of a moderating influence of hippocampal volume on the strength of associations between its functional activity and cognitive performance (e.g., Hou et al.,2020), are indicative of a complex relationship between a neural region’s size and its functional capacity (Mooraj et al., 2025). In short, null associations between the volume of a given hippocampal subfield and performance on a cognitive task should not be taken as evidence that the subfield makes no contribution to the computations that support task performance (see also Clark et al., 2023), a point to be borne in mind when considering the present (and prior) findings.

The present study suffers from other limitations also. One is its cross-sectional design, meaning that age effects cannot be attributed unambiguously to aging rather than to confounding factors such as selection bias or cohort effects (Rugg, 2016). A second limitation concerns the employment of a semi-automated segmentation procedure which, while demonstrating good agreement with manual segmentation (Yushkevich et al., 2015a), likely introduced measurement noise into the delineation of the different hippocampal subfields. Third, our study sample was predominantly well educated and of relatively high socio-economic status, potentially limiting the generality of the findings. Lastly, the study was only moderately powered, and hence unable to detect small effects. Thus, it remains to be determined which, if any, of the smaller effects reported in Tables 7 and 8 that failed to survive correction for multiple comparisons are reliable.

## Acknowledgements

We are grateful to all of our participants for volunteering their time for this study.

## References

Aghjayan, S. L., Polk, S. E., Ripperger, H. S., Huang, H., Wan, L., Kamarck, T., Marsland, A. L., Kang, C., Voss, M. W., Sutton, B. P., Oberlin, L. E., Burns, J. M., Vidoni, E. D., McAuley, E., Hillman, C. H., Kramer, A. F., & Erickson, K. I. (2025). Associations Between Episodic Memory and Hippocampal Volume in Late Adulthood. Hippocampus, 35(2), e70010-n/a. 10.1002/hipo.70010

Amat, J. A., Bansal, R., Whiteman, R., Haggerty, R., Royal, J., & Peterson, B. S. (2008). Correlates of intellectual ability with morphology of the hippocampus and amygdala in healthy adults. Brain and Cognition, 66(2), 105–114. 10.1016/j.bandc.2007.05.009

Aribisala, B. S., Royle, N. A., Maniega, S. M., Valdés Hernández, M. C., Murray, C., Penke, L., Gow, A., Starr, J. M., Bastin, M. E., Deary, I. J., & Wardlaw, J. M. (2014). Quantitative multi-modal MRI of the Hippocampus and cognitive ability in community-dwelling older subjects. Cortex, 53(100), 34–44. 10.1016/j.cortex.2013.12.012

Aslaksen, P. M., Bystad, M. K., Ørbo, M. C., & Vangberg, T. R. (2018). The relation of hippocampal subfield volumes to verbal episodic memory measured by the California Verbal Learning Test II in healthy adults. Behavioural Brain Research, 351, 131–137. 10.1016/j.bbr.2018.06.008

Aumont, E., Bussy, A., Bedard, M.-A., Bezgin, G., Therriault, J., Savard, M., Fernandez Arias, J., Sziklas, V., Vitali, P., Poltronetti, N. M., Pallen, V., Thomas, E., Gauthier, S., Kobayashi, E., Rahmouni, N., Stevenson, J., Tissot, C., Chakravarty, M. M., & Rosa-Neto, P. (2023). Hippocampal subfield associations with memory depend on stimulus modality and retrieval mode. Brain Communications, 5(6), fcad309. 10.1093/braincomms/fcad309

Benedict, R. H. B. (1997). Brief visuospatial memory test–revised: Professional manual. Lutz, FL: Psychological Assessment Resources.

Benton, A. L. (1968). Differential behavioral effects in frontal lobe disease. Neuropsychologia, 6(1), 53–60.

Botdorf, M., Canada, K. L., & Riggins, T. (2022). A meta-analysis of the relation between hippocampal volume and memory ability in typically developing children and adolescents. Hippocampus, 32(5), 386–400. 10.1002/hipo.23414

Canada, K. L., Mazloum-Farzaghi, N., Rådman, G., Adams, J. N., Bakker, A., Baumeister, H., Berron, D., Bocchetta, M., Carr, V. A., Dalton, M. A., Flores, R., Keresztes, A., La Joie, R., Mueller, S. G., Raz, N., Santini, T., Shaw, T., Stark, C. E. L., Tran, T. T., … Daugherty, A. M. (2024). A (sub)field guide to quality control in hippocampal subfield segmentation on high-resolution T2-weighted MRI. Human Brain Mapping, 45(15), e70004-n/a. 10.1002/hbm.70004

Cipolotti, L., Ruffle, J. K., Mole, J., Xu, T., Hyare, H., Shallice, T., Chan, E., & Nachev, P. (2023). Graph lesion-deficit mapping of fluid intelligence. Brain (London, England : 1878), 146(1), 167–181. 10.1093/brain/awac304

Clark, I. A., Monk, A. M., Hotchin, V., Pizzamiglio, G., Liefgreen, A. (2020). Does hippocampal volume explain performance differences on hippocampal-dependent tasks? Neuroimage 221, 117211. 10.1016/j.neuroimage.2020.117211

Clark, I. A., Dalton, M. A., and Maguire, E. A. (2023). Posterior hippocampal CA2/3 volume is associated with autobiographical memory recall ability in lower performing individuals. Scientific Reports, 13:7924. 10.1038/s41598-023-35127-2

Colom, R., Stein, J. L., Rajagopalan, P., Martínez, K., Hermel, D., Wang, Y., Álvarez-Linera, J., Burgaleta, M., Quiroga, M. Á., Shih, P. C., & Thompson, P. M. (2013). Hippocampal structure and human cognition: Key role of spatial processing and evidence supporting the efficiency hypothesis in females. Intelligence (Norwood), 41(2), 129–140. 10.1016/j.intell.2013.01.002

Cooper, E., Greve, A., & Henson, R. N. (2017). Assumptions behind scoring source versus item memory: Effects of age, hippocampal lesions and mild memory problems. Cortex, 91, 297–315. 10.1016/j.cortex.2017.01.001

Daugherty, A. M., Bender, A. R., Raz, N., & Ofen, N. (2016). Age differences in hippocampal subfield volumes from childhood to late adulthood. Hippocampus, 26(2), 220–228. 10.1002/hipo.22517

de Chastelaine, M., Srokova, S., Hou, M., Kidwai, A., Kafafi, S. S., Racenstein, M. L., & Rugg, M. D. (2023). Cortical thickness, gray matter volume, and cognitive performance: a cross-sectional study of the moderating effects of age on their interrelationships. Cerebral Cortex, 33(10), 6474–6485. 10.1093/cercor/bhac518

Delis, D. C., Kramer, J. H., Kaplan, E., & Ober, B. A. (2000). California verbal learning test (2nd ed.). San Antonio, TX: Psychological Corporation.

Fjell, A. M., Sneve, M. H., Amlien, I. K., Grydeland, H., Mowinckel, A. M., Vidal-Piñeiro, D., Sørensen, Ø., & Walhovd, K. B. (2025). Stable hippocampal correlates of high episodic memory function across adulthood. Scientific Reports, 15(1), Article 8816. 10.1038/s41598-025-92278-0

Foster, C. M., Kennedy, K. M., Hoagey, D. A., & Rodrigue, K. M. (2019). The role of hippocampal subfield volume and fornix microstructure in episodic memory across the lifespan. Hippocampus, 29(12), 1206–1223. 10.1002/hipo.23133

Gohel, S., Chan, C. C., Baptista, I., & Szeszko, P. R. (2025). Age-associated trajectories of hippocampus subregion volumes from childhood through later adulthood. Neurobiology of Aging, 156, 73–84. 10.1016/j.neurobiolaging.2025.08.002

Henson, R.N., Campbell, K.L., Davis, S.W., Taylor, J.R., Emery, T., Erzinclioglu, S.; Cam-CAN; Kievit, R.A. (2016). Multiple determinants of lifespan memory differences. Scientific Reports, 7;6:32527. 10.1038/srep32527

Hindy, N. C., Ng, F. Y., & Turk-Browne, N. B. (2016). Linking pattern completion in the hippocampus to predictive coding in visual cortex. Nature Neuroscience, 19(5), 665–667. 10.1038/nn.4284

Homayouni, R., Canada, K. L., Saifullah, S., Foster, D. J., Thill, C., Raz, N., Daugherty, A. M., & Ofen, N. (2023). Age-related differences in hippocampal subfield volumes across the human lifespan: A meta-analysis. Hippocampus, 33(12), 1292–1315. 10.1002/hipo.23582

Hou, M., de Chastelaine, M., Donley, B. E., & Rugg, M. D. (2021). Specific and general relationships between cortical thickness and cognition in older adults: a longitudinal study. Neurobiology of Aging, 102, 89–101. 10.1016/j.neurobiolaging.2020.11.004

Hou, M., de Chastelaine, M., Jayakumar, M., Donley, B. E., & Rugg, M. D. (2020). Recollection-related hippocampal fMRI effects predict longitudinal memory change in healthy older adults. Neuropsychologia, 146, 107537. 10.1016/j.neuropsychologia.2020.107537

Iglesias, J. E., Augustinack, J. C., Nguyen, K., Player, C. M., Player, A., Wright, M., Roy, N., Frosch, M. P., McKee, A. C., Wald, L. L., Fischl, B., & Van Leemput, K. (2015). A computational atlas of the hippocampal formation using ex vivo, ultra-high resolution MRI: Application to adaptive segmentation of in vivo MRI. *NeuroImage* (Orlando, Fla.), 115, 117–137. 10.1016/j.neuroimage.2015.04.042

Lai, Y.-M., & Chang, Y.-L. (2023). Age-related differences in associative memory recognition of Chinese characters and hippocampal subfield volumes. Biological Psychology, 183, Article 108657. 10.1016/j.biopsycho.2023.108657

Mattson J. T., Wang T. H., de Chastelaine M., Rugg M. D. (2014). Effects of age on negative subsequent memory effects associated with the encoding of item and item-context information. Cereb Cortex, 24(12):3322–3333. 10.1093/cercor/bht193

Miller, T. D., Chong, T. T., Aimola Davies, A. M. Ng, T. W. C., Johnson, M. R., Irani, S. R., Vincent, A., Husain, M., Jacob, S., Maddison, P., Kennard, C., Gowland, P. A., Rosenthal, C. R. (2017). Focal CA3 hippocampal subfield atrophy following LGI1 VGKC-complex antibody limbic encephalitis. Brain, 140, 1212–1219. 10.1093/brain/awx070

Mooraj, Z., Salami, A., Campbell, K. L., Dahl, M. J., Kosciessa, J. Q., Nassar, M. R., Werkle-Bergner, M., Craik, F. I. M., Lindenberger, U., Mayr, U., Rajah, M. N., Raz, N., Nyberg, L., & Garrett, D. D. (2025). Toward a functional future for the cognitive neuroscience of human aging. Neuron, 113(1), 154–183. 10.1016/j.neuron.2024.12.008

O’Reilly, R. C., Bhattacharyya, R., Howard, M. D., & Ketz, N. (2014). Complementary Learning Systems. Cognitive Science, 38(6), 1229–1248. 10.1111/j.1551-6709.2011.01214.x

O’Shea, A., Cohen, R. A., Porges, E. C., Nissim, N. R., & Woods, A. J. (2016). Cognitive Aging and the Hippocampus in Older Adults. Frontiers in Aging Neuroscience, 8, 298. 10.3389/fnagi.2016.00298

Oechslin, M. S., Descloux, C., Croquelois, A., Chanal, J., Van De Ville, D., Lazeyras, F., & James, C. E. (2013). Hippocampal volume predicts fluid intelligence in musically trained people. Hippocampus, 23(7), 552–558. 10.1002/hipo.22120

Pohlack, S. T., Meyer, P., Cacciaglia, R., Liebscher, C., Ridder, S., & Flor, H. (2014). Bigger is better! Hippocampal volume and declarative memory performance in healthy young men. Brain Structure and Function, 219(1), 255–267. 10.1007/s00429-012-0497-z

Raven J., Raven J. C., Courth J. H. (2000). Manual for Raven’s progressive matrices and vocabulary scales. Section 4: the advanced progressive matrices. San Antonio, TX: Harcourt Assessment.

Raz, N., Lindenberger, U., Ghisletta, P., Rodrigue, K. M., Kennedy, K. M., & Acker, J. D. (2008). Neuroanatomical Correlates of Fluid Intelligence in Healthy Adults and Persons with Vascular Risk Factors. Cerebral Cortex (New York, N.Y. 1991), 18(3), 718–726. 10.1093/cercor/bhm108

Reitan, R. M., & Wolfson, D. (1985). The Halstead–Reitan neuropsychological test battery: Therapy and clinical interpretation. Tucson, AZ: Neuropsychological Press.

Rempel-Clower, N. L., Zola, S. M., Squire, L. R., & Amaral, D. G. (1996). Three Cases of Enduring Memory Impairment after Bilateral Damage Limited to the Hippocampal Formation. The Journal of Neuroscience, 16(16), 5233–5255. 10.1523/jneurosci.16-16-05233.1996

Reuben, A., Brickman, A. M., Muraskin, J., Steffener, J., & Stern, Y. (2011). Hippocampal Atrophy Relates to Fluid Intelligence Decline in the Elderly. Journal of the International Neuropsychological Society, 17(1), 56–61. 10.1017/S135561771000127X

Rugg, M. D. (2016). Interpreting Age-Related Differences in Memory-Related Neural Activity. In Cognitive Neuroscience of Aging (pp. 183–204). Oxford University Press. 10.1093/acprof:oso/9780199372935.003.0008

Rugg, M. D., & Vilberg, K. L. (2013). Brain networks underlying episodic memory retrieval. Current opinion in neurobiology, 23(2), 255–260. 10.1016/j.conb.2012.11.005

Scoville, W. B., & Milner, B. (1957). Loss of recent memory after bilateral hippocampal lesions. *Journal of Neurology*, Neurosurgery and Psychiatry, 20(1), 11–21. 10.1136/jnnp.20.1.11

Shing, Y. L., Rodrigue, K. M., Kennedy, K. M., Fandakova, Y., Bodammer, N., Werkle-Bergner, M., Lindenberger, U., & Raz, N. (2011). Hippocampal Subfield Volumes: Age, Vascular Risk, and Correlation with Associative Memory. Frontiers in Aging Neuroscience, 3, 2. 10.3389/fnagi.2011.00002

Singh, D., & Schapiro, A. C. (2026). Evidence for complementary learning systems within the hippocampus. Philosophical transactions of the Royal Society of London. Series B, Biological sciences, 381(1954), 20250243. 10.1098/rstb.2025.0243

Smith, A. (1973). Symbol digit modalities test. Los Angeles: Western Psychological Services.

Snodgrass JG, Corwin J (1988). Pragmatics of Measuring Recognition Memory: Applications to Dementia and Amnesia. J Exp Psychol Gen. 1988;117(1):34–50.

Spreen, O., & Benton, A. L. (1977). Neurosensory center comprehensive examination for aphasia. Victoria, Canada: Neuropsychology Laboratory.

Tenderra, R. M., & Theves, S. (2025). Human intelligence relates to neural measures of cognitive map formation. Cell Reports (Cambridge), 44(8), Article 116033. 10.1016/j.celrep.2025.116033

Van Petten, C. (2004). Review of Relationship between hippocampal volume and memory ability in healthy individuals across the lifespan: review and meta-analysis. Neuropsychologia, 42(10), 1394–1413. 10.1016/j.neuropsychologia.2004.04.006

Wechsler, D. (1981). WAIS-R: Wechsler Adult Intelligence Scale-Revised. New York: Psychological Corporation.

Wechsler, D. (2009). Wechsler Memory Scale (4th ed.). San Antonio, TX: Psychological Corporation.

Wechsler D. 2011. Test of premorbid functioning. San Antonio, TX: The Psychological Corporation.

Winterburn, J. L., Pruessner, J. C., Chavez, S., Schira, M. M., Lobaugh, N. J., Voineskos, A. N., & Chakravarty, M. M. (2013). A novel in vivo atlas of human hippocampal subfields using high-resolution 3 T magnetic resonance imaging. NeuroImage (Orlando, Fla.), 74, 254–265. 10.1016/j.neuroimage.2013.02.003

Yang, Y., Wang, D., Hou, W., Li, H. (2023). Cognitive Decline Associated with Aging. In: Zhang, Z. (eds) Cognitive Aging and Brain Health. Advances in Experimental Medicine and Biology, vol 1419. Springer, Singapore. 10.1007/978-981-99-1627-6_3

Yassa, M. A., & Stark, C. E. L. (2011). Pattern separation in the hippocampus. Trends in Neurosciences (Regular Ed.), 34(10), 515–525. 10.1016/j.tins.2011.06.006

Yonelinas, A. P., Kroll, N. E. A., Dobbins, I., Lazzara, M., & Knight, R. T. (1998). Recollection and Familiarity Deficits in Amnesia: Convergence of Remember-Know, Process Dissociation, and Receiver Operating Characteristic Data. Neuropsychology, 12(3), 323–339. 10.1037/0894-4105.12.3.323

