## Supplemental Tables 1-6 for "Different hippocampal subfield volumes predict source memory performance and general cognitive ability in an adult lifespan sample"

**Supplemental Table 1**. Results of one-way ANCOVAs (covariates of sex and years of education) examining the effects of age group (young, middle-aged, older) on each cognitive measure and *post-hoc* pairwise contrasts.

| | |
|---|---|
| *Speed* | |
| Age group | $F(2,158) = 42.74$, $p < .001$, partial $\eta^2 = .351$ |
| *Pairwise-comparison* | |
| Young vs middle | $p = .533$ |
| Young > older | $p < .001$ |
| Middle > older | $p < .001$ |
| *Memory 1* | |
| Age group | $F(2,158) = 19.43$, $p < .001$, partial $\eta^2 = .197$ |
| *Pairwise-comparison* | |
| Young > middle | $p = .002$ |
| Young > older | $p < .001$ |
| Middle > older | $p = .006$ |
| *Fluency* | |
| Age group | $F(2,158) = 9.17$, $p < .001$, partial $\eta^2 = .104$ |
| *Pairwise-comparison* | |
| Young vs middle | $p = .352$ |
| Young > older | $p = .010$ |
| Middle > older | $p < .001$ |

| | |
|---|---|
| <u>*Memory 2*</u> | |
| Age group | $F(2,158) = 23.26$, $p < .001$, partial $\eta^2 = .227$ |
| *Pairwise-comparison* | |
| Young > middle | $p = .004$ |
| Young > older | $p < .001$ |
| Middle > older | $p < .001$ |
| <u>*Span*</u> | |
| Age group | $F(2,158) = 4.533$, $p = .012$, partial $\eta^2 = .054$ |
| *Pairwise-comparison* | |
| Young vs middle | $p = .327$ |
| Young vs older | $p = .120$ |
| Middle > older | $p = .004$ |
| <u>*General Cognitive Ability*</u> | |
| Age group | $F(2,158) = 28.959$, $p < .001$, partial $\eta^2 = .268$ |
| *Pairwise-comparison* | |
| Young vs middle | $p = .228$ |
| Young > older | $p < .001$ |
| Middle > older | $p < .001$ |
| <u>*pSR*</u> | |
| Age group | $F(2,158) = 11.306$, $p < .001$, partial $\eta^2 = .125$ |
| *Pairwise-comparison* | |
| Young > middle | $p = .005$ |
| Young > older | $p < .001$ |
| Middle vs older | $p = .102$ |

| | |
|---|---|
| *Pr* | |
| Age group | F (2,158) = 18.668, $p < .001$, partial $\eta^2 = .191$ |
| *Pairwise-comparison* | |
| Young > middle | $p = .007$ |
| Young > older | $p < .001$ |
| Middle > older | $p = .002$ |

**Supplemental Table 2**. Results of one-way ANCOVAs (covariates of sex and years of education) examining the effects of age group (young, middle-aged, older) on each hippocampal subfield and *post-hoc* pairwise contrasts.

| | |
|---|---|
| *CA1* | |
| Age group | F (2,158) = 4.749, p =.010, partial $\eta^2$ = .057 |
| Pairwise-comparison | |
| Young vs middle | p = .387 |
| Young vs older | p = .087 |
| Middle > older | p = .004 |
| *CA2-3* | |
| Age group | F (2,158) = 1.651, p =.195, partial $\eta^2$ = .020 |
| *DG* | |
| Age group | F (2,158) = 0.890, p =.413, partial $\eta^2$ = .011 |
| *SUBICULUM* | |
| Age group | F (2,158) = 5.416, p =.005, partial $\eta^2$ = .064 |
| Pairwise-comparison | |
| Young < middle | p = .033 |
| Young vs older | p = .592 |
| Middle > older | p = .001 |

**Supplemental Table 3**. Hierarchical multiple regression results for dentate gyrus volume predicting non-mnemonic composite component score.

| | Standardized β | t (p) | F (p) | ΔF (p) | $R^2$ | $\Delta R^2$ |
|---|---|---|---|---|---|---|
| *Step 1* | | | 13.72 (< .001) | | 0.26 | |
| Intercept | | -1.28 (.202) | | | | |
| Sex | 0.00 | 0.05 (.960) | | | | |
| Years of education | 0.14 | 1.89 (.061) | | | | |
| Linear age | -0.64 | -7.37 (< .001) | | | | |
| Quadratic age | -0.35 | -4.07 (< .001) | | | | |
| *Step 2* | | | 13.23 (< .001) | 8.63 (.004) | 0.30 | 0.04 |
| DG | 0.20 | 2.94 (.004) | | | | |

**Supplemental Table 4**. Multiple regression results for sex, years of education and age predicting whole hippocampal volume.

| | Standardized β | t (p) | F (p) | $R^2$ |
|---|---|---|---|---|
| *Whole hippocampus* | | | 2.22 (.070) | 0.05 |
| Intercept | | -0.14 (.892) | | |
| Sex | -0.02 | -0.28 (.782) | | |
| Years of education | 0.05 | 0.58 (.563) | | |
| Linear age | -0.26 | -2.64 (.009) | | |
| Quadratic Age | -0.25 | -2.52 (.013) | | |

**Supplemental Table 5**. Hierarchical multiple regression results for whole hippocampal volume predicting pSR.

| Parameter | Standardized β | t (p) | F (p) | ΔF (p) | $R^2$ | $\Delta R^2$ |
|---|---|---|---|---|---|---|
| *Step 1* | | | 6.82 (< .001) | | 0.15 | |
| Intercept | | 3.42 (< .001) | | | | |
| Sex | 0.00 | -0.00 (.997) | | | | |
| Years of education | 0.05 | 0.63 (.531) | | | | |
| Linear age | -0.40 | -4.28 (< .001) | | | | |
| Quadratic age | -0.10 | -0.10 (.918) | | | | |
| *Step 2* | | | 6.93 (< .001) | 6.43 (.012) | 0.18 | 0.34 |
| Whole hippocampus | 0.19 | 2.54 (.012) | | | | |

**Supplemental Table 6**. Hierarchical multiple regression results for whole hippocampal volume predicting general cognitive ability.

| Parameter | Standardized β | t (p) | F (p) | ΔF (p) | $R^2$ | $\Delta R^2$ |
|---|---|---|---|---|---|---|
| _Step 1_ | | | 20.41 (< .001) | | 0.34 | |
| Intercept | | -2.05 (.042) | | | | |
| Sex | -0.13 | -2.02 (.045) | | | | |
| Years of education | 0.19 | 2.84 (.005) | | | | |
| Linear age | -0.70 | -8.53 (< .001) | | | | |
| Quadratic age | -0.27 | -3.34 (.001) | | | | |
| _Step 2_ | | | 18.31 (< .001) | 6.86 (.010) | 0.37 | 0.03 |
| Whole hippocampus | 0.17 | 2.62 (.010) | | | | |
